# T cells Use Focal Adhesions to Pull Themselves Through Confined Environments

**DOI:** 10.1101/2023.10.16.562587

**Authors:** Alexia Caillier, David Oleksyn, Deborah J. Fowell, Jim Miller, Patrick W. Oakes

## Abstract

Immune cells are highly dynamic and able to migrate through environments with diverse biochemical and mechanical composition. Their migration has classically been defined as amoeboid under the assumption that it is integrin-independent. Here we show that activated primary Th1 T cells require both confinement and extracellular matrix protein to migrate efficiently. This migration is mediated through small and dynamic focal adhesions that are composed of the same proteins associated with canonical mesenchymal focal adhesions, such as integrins, talin, and vinculin. These focal adhesions, furthermore, localize to sites of contractile traction stresses, enabling T cells to pull themselves through confined spaces. Finally, we show that Th1 T cell preferentially follows tracks of other T cells, suggesting that these adhesions are modifying the extracellular matrix to provide additional environmental guidance cues. These results demonstrate not only that the boundaries between amoeboid and mesenchymal migration modes are ambiguous, but that integrin-mediated adhesions play a key role in T cell motility.

**Graphical Abstract:** 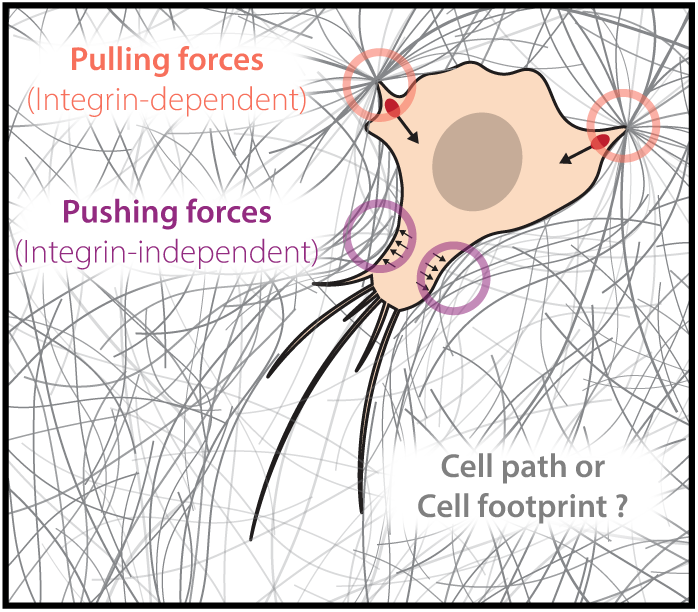

## Introduction

Migration is a critical component of an efficient immune response. Cells must be ready to respond to diverse signalling cascades, and then physically move to various sites throughout the body in response to those signals. While biochemical signals such as chemokines have long been associated with stimulating and facilitating migration, more recent studies have implicated both passive and active biophysical signals as playing equally integral roles [1–4]. Indeed, passive features of the extracellular environment including it’s stiffness [5–8], composition [9, 10], and architecture [11–13] can all influence cell migration [1, 14]. Active forces generated internally by the cell and by surrounding cells in the tissue can also have an impact [15–18], such as by modulating bond strength and lifetime. T cells, which must navigate a vast environmental complexity - from the lymphatic system, to the circulatory system, and then the surrounding tissues - are particularly well positioned to display adaptive migration mechanisms depending on their given environment.

Much of our foundational knowledge of the molecular mechanisms of migration are derived from studying mesenchymal cells on account of their large, flat morphology and reasonably slow dynamics [19]. This combination makes them ideal for high-resolution microscopy. Mesenchymal migration is characterized by a narrow band of actin polymerization at a leading edge, the formation of large integrin mediated focal adhesion (FA) plaques, and bundled actomyosin stress fibers in the cell body [20]. In contrast, immune cells have typically been described as navigating their complex environment using an amoeboid mode of migration [21]. Amoeboid migration is characterized by a more rounded cell morphology and a dependence on rapid bulk actin polymerization [22]. While the term amoeboid was first used to describe migration in amoeba like Dictyostelium, it has generally evolved to act as a catch-all for migration that is not integrin mediated [22]. The molecular mechanisms mediating this form of migration have remained unclear, but are generally ascribed to non-specific adhesion and friction forces between the cell and the surrounding environment, enabled by internal fluid flows [23–26].

T cells and other leukocytes express a number of different integrins, including those that bind extracellular matrix (ECM) proteins like Intercellular Adhesion Molecule 1 (ICAM-1), fibronectin (FN) and collagen [27–29]. This diverse array of binding proteins should be useful for navigating the many complex extracellular environments encountered during an immune response. It was thus surprising when it was reported that dendritic cells could migrate effectively in 3D geometries when all integrins were knocked out [29]. It has been further suggested that integrin-independent migration enables immune cells to migrate through tissue and interstitial spaces after crossing the endothelial layer [30–32]. More recently, however, it was shown that not all immune cells migrate integrin-independently [33, 34]. Integrin-collagen interactions were previously shown to be essential for efficient immune response in several mouse models [35, 36]. It was also shown that helper T cells overexpress fibronectin-specific integrins following their activation, and that those integrins are required for migration in vivo [28]. More specifically, CD4+ T cell interstitial migration is highly dependent on α*_v_* integrin adhesion with ECM [37, 38]. Furthermore, α*_v_*β_3_ integrins were very recently shown to be essential for efficient adaptative helper T cell immune response, as the inhibition of α*_v_*β_3_ completely impaired adaptative immune response [39]. The contrast in these findings highlights the need to understand the molecular mechanisms that underlie immune cell migration.

In this paper we demonstrate that primary Th1 T cells form FAs which they use to migrate, and that FA formation depends on the extracellular environment composition and geometry. We find that Th1 cells require integrin mediated adhesion to migrate, and in the case of the FN, cells additionally require confinement to move efficiently. When Th1 cells make FAs, they are consistent in composition to canonical mesenchymal FAs, and are sites where cytoskeletal contractile forces are transmitted to the ECM. These findings challenge the traditional framing of T cell migration as both amoeboidal and integrin-independent. Finally we show that T cells tend to follow in the tracks of other T cells, suggesting additional functional roles for FAs in enabling a rapid and directed multicellular immune response.

## Results

### Th1 migration is regulated by environment composition and geometry

Using activated primary Th1 cells isolated from OTII mice (Fig. S1A-F), we first tested whether T cells could migrate on glass substrates coated with either FN (likely to be found in the tissue ECM and a ligand for both α*_v_*β_3_ and α_5_β_1_) or ICAM-1 (found on the surface of other cells and a ligand for α*_L_*β_2_). As expected, we found that Th1 cells migrated fast and efficiently on ICAM-1 (Fig. 1A-B). On FN, in contrast, cells exhibited Brownian motion and no migration, only transiently interacting with the coverslip surface (Fig. 1A-B). When confined under a PDMS surface with a gap of 5µm, however, Th1 cells migrated robustly on both FN and ICAM-1 with speeds and displacements that were indistinguishable from unconfined Th1 cells migrating on ICAM-1 (Fig. 1A-B).

**Fig. 1.**
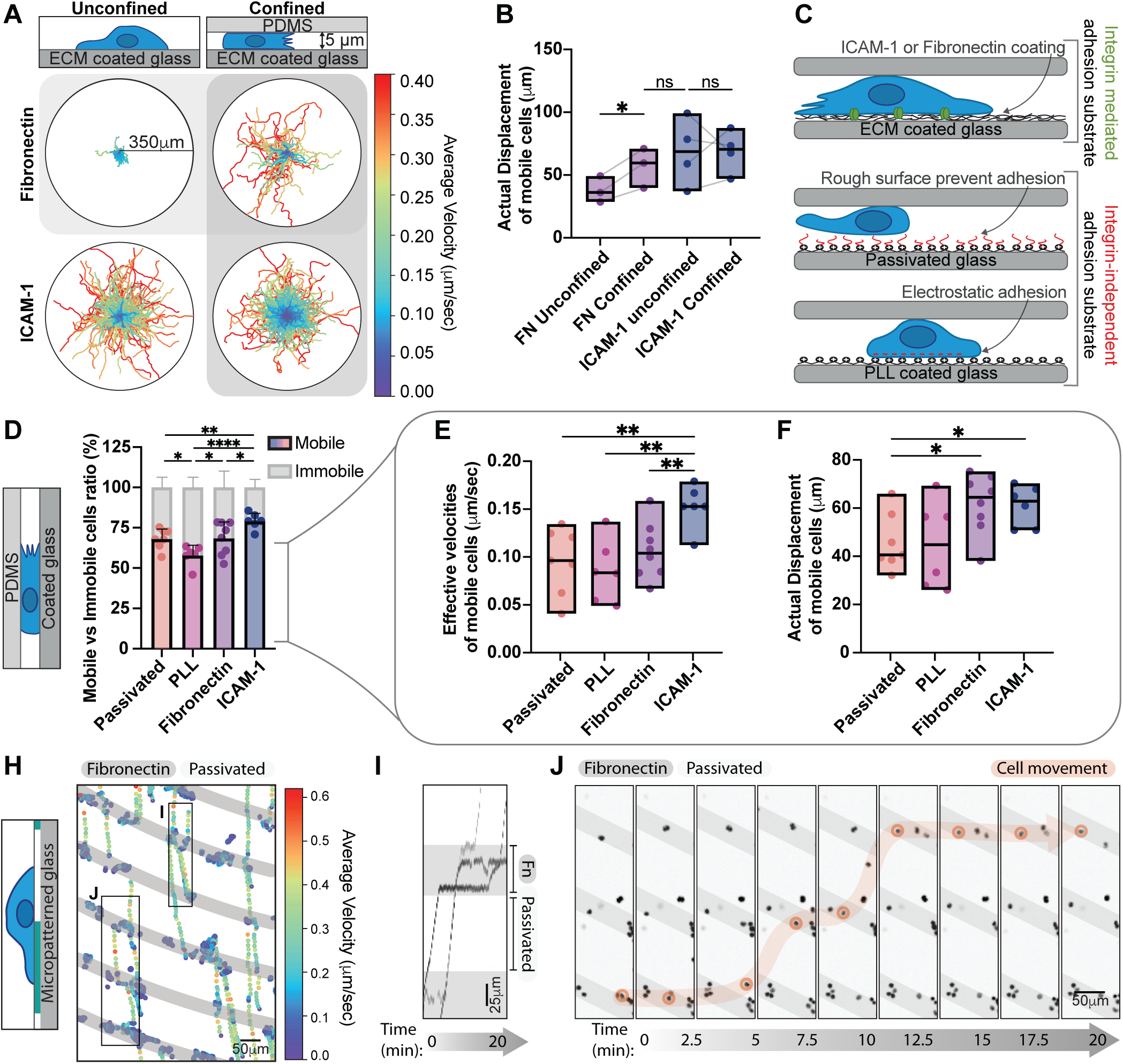
Th1 migration, velocity and displacement are regulated by the environment composition and geometry. (A) Th1 cell tracking on ICAM-1 and Fibronectin in unconfined vs confined (5*µ*m) environments. The colormap represents the cell average velocity in *µ*m/sec. (B) Comparison of the actual displacement, defined as the distance between starting and end points of a track, of the mobile cells in (A) . (C) Graphical representation of the different substrates used and their properties. (D) The mobile versus immobile fraction of cells in confinement on ICAM-1, Fibronectin, PLL and Passivated surfaces. Mobile cells are defined by a minimum displacement of 10*µ*m. The effective velocities (E) and actual displacement (F) of mobile cells in (D). (H) Cell tracking on a FN micropatterned substrate. Grey = FN, White = Passivated. The colors represent the instantaneous velocities of each cell at a give time point. (I) Kymographs of the regions identified in (H) showing cells stalling on the FN patterned stripes. (J) Snapshots of the movie from (H). A single cell is highlighted in pink to illustrate its trajectory over time.

To explore this further, we compared integrin-mediated migration of Th1 cells confined on either FN or ICAM-1, with integrin independent migration of Th1 cells confined on surfaces coated either with electrostatically charged polymers (Poly-L-lysine; PLL), or inert blocking polymers (PLL-PEG or PMOXA; Passivated)(Fig. 1C). We found that over 75% of cells were mobile on ICAM-1, while only approximately 50% were mobile on PLL, with cells on FN or passivated surfaces in between (Fig. 1D, Supplementary Video 1). If we compared the effective velocity and displacement of the mobile fraction, cells plated on integrin mediated surfaces moved larger distances than those on integrin-independent surfaces, and were fastest on ICAM-1 (Fig. 1E-F, Supplementary Video 1).

To confirm that Th1 cells were directly interacting with FN we used micropatterning to create stripes of fibronectin in an otherwise passivated surface (Fig. 1H). Convective flows induced by imaging multiple positions allowed us to track the motion of unconfined cells as they crossed different regions. We found that Th1 cells slowed down and clustered along the FN coated regions (Fig. 1H, Supplementary Video 2), lingering for brief periods of time before detaching and moving on to the next region (Fig. 1I-J; Supplementary Video 2). Together these results indicate that Th1 cells are able to migrate effectively on FN when under confinement and that this migration is mediated through specific interaction with FN.

### ECM is necessary for Th1 migration

Our initial experiments suggested that Th1 T cells could engage both integrin mediated and integrin-independent migration modes, although with varying degrees of efficacy, when confined on different substrates. Fully passivating a surface is challenging, however, and previous results have shown that these surfaces are still able to adsorb many proteins present in serum, such as ECM [40]. To ensure that the ECM present in cell culture media serum [41, 42] doesn’t bind to our passivated surfaces and contribute to Th1 migration, we repeated our confined migration experiments, switching the cells to serum free media immediately prior to confinement (Fig. 2A). Surprisingly, the mobile fraction dropped precipitously for cells on PLL or passivated substrates (Fig. 2A,D) and cells exhibited significantly reduced velocities and displacements (Fig. 2B-C). Th1 cells confined on both 10 µg/mL FN and ICAM-1 in serum free media, however, exhibited similar mobile fractions (Fig. 2A vs Fig. 1D), velocities and displacements as cells plated in full serum media (Figs. 2B-D vs 1E-F and Fig. S1G-H, Supplementary Video 3).

**Fig. 2.**
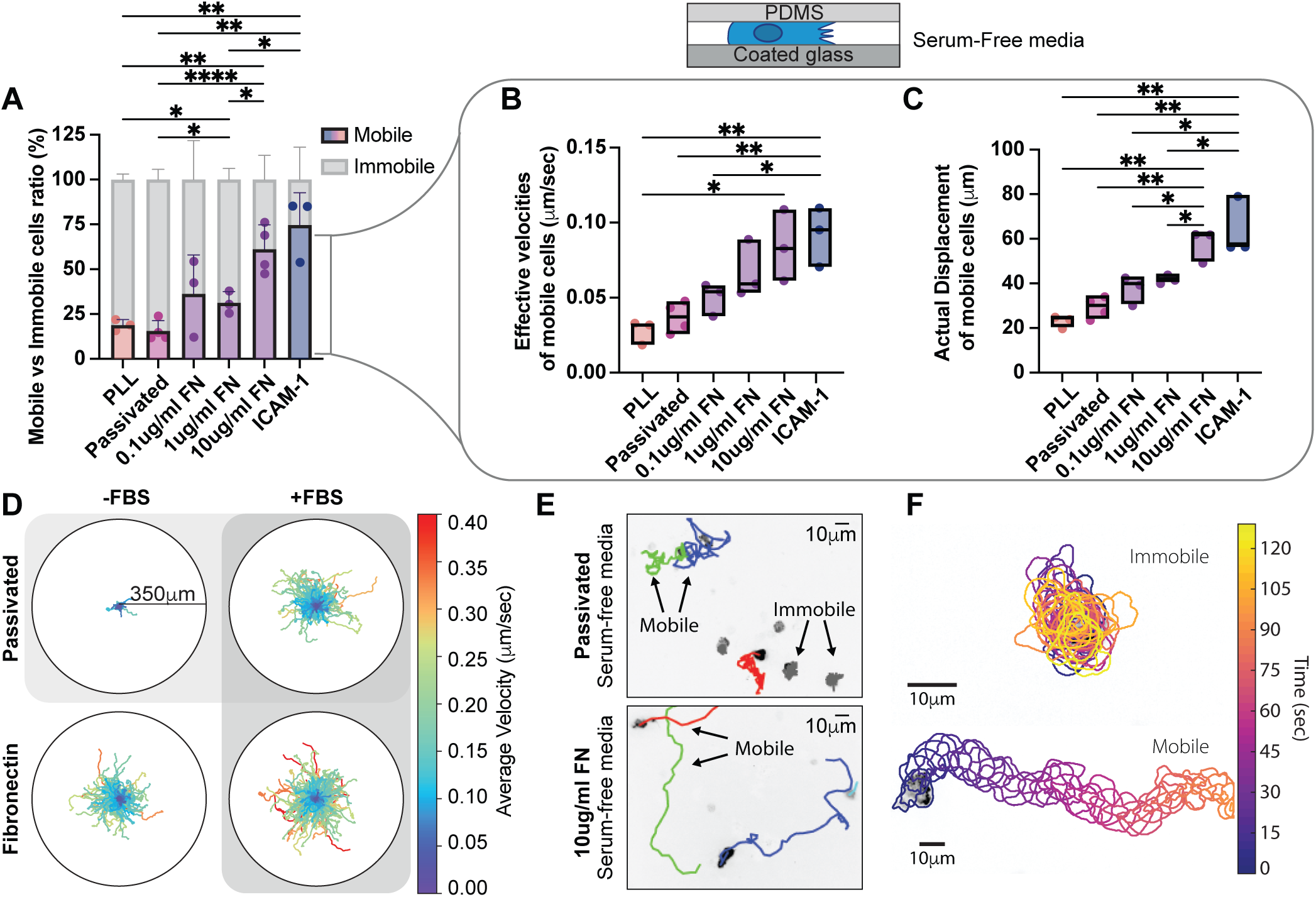
ECM is essential for Th1 migration. (A) The mobile versus immobile Th1 cells in confinement on ICAM-1, Fibronectin, PLL and Passivated surface in serum-free media. Mobile cells are defined by a minimum displacement of 10 *µ*m. Analysis of the effective velocities (B) and the actual displacement (C) of mobile cells in (A). (D) Cell tracks on passivated substrates versus on FN in either serum-free media (-FBS) or with serum (+FBS). The colors represent the cell’s average velocity. (E) Tracking of cells on passivated versus FN substrates in serum-free media. (F) Representative immobile and mobile cells migrating in serum-free media. The outlines are color coded for time.

Moreover, this effect was titratable, as the mobile fraction, velocity and displacement, were proportional to the concentration of FN (Fig. 2A-C). High resolution imaging of cells migrating in serum-free media on passivated surface presented active membrane ruffles and protrusions indicating that cells were attempting and failing to move on the integrin-independent surface, remaining active with short term serum depletion (Fig. 2E-F; Supplementary Video 4). Cells migrating on fibronectin coated surfaces in serum-free media also presented active membrane ruffles and protrusions (Fig. 2E-F; Supplementary Video 4), but the absence of serum had no significant impact on their migration in speed or displacement (Fig. S1G-H). To confirm that confinement alone impacted Th1 migration on FN, we repeated the experiment in Fig. 1B, but in serum-free media (Fig.S 1I). Th1 cells in serum-free media still migrated further and faster on FN in confinement then when unconfined. Together these results suggest that integrin-dependant substrates are necessary for robust Th1 migration.

### Th1 cells form focal adhesions

The necessity of integrin dependent substrates for migration suggests that Th1 T cells are forming canonical integrin-based adhesions. Using RNA-seq data from a publicly available database (Th-express.org [43]), we observe that activated Th1 express RNA for canonical FA proteins (e.g. talin, paxillin, zyxin, vinculin), in addition to the integrins required to bind ECM (Fig. 3A). Western blots of lysates from activated Th1 corroborate the presence of FA components at the protein level (Fig. S2F) To determine whether these proteins were coalescing into actual FA, we imaged Th1 cells transduced with either talin-EGFP (Fig. 3E-H, Supplementary Video 5) or vinculin-EGFP (Fig. S2A-D). On both ICAM-1 (Fig. 3E and S2A) and on Fibronectin (Fig 3F and S2B) we observed highly dynamic FA-like structures (Supplementary Video 5), whereas we were unable to observe any such structures on PLL (Fig. 3G and S2C, Supplementary Video 5) or passivated substrates (Figs. 3H and S2D, Supplementary Video 5). As a comparison, we measured FA lifetime in Th1 cells compared to the same constructs in a mouse fibroblast cell line. We found that the FAs in Th1 T cells were significantly shorter-lived compared to the fibroblast, with an average lifetime of only 1.7 minutes compared to 40 min in the fibroblasts (Fig. 3B).

**Fig. 3.**
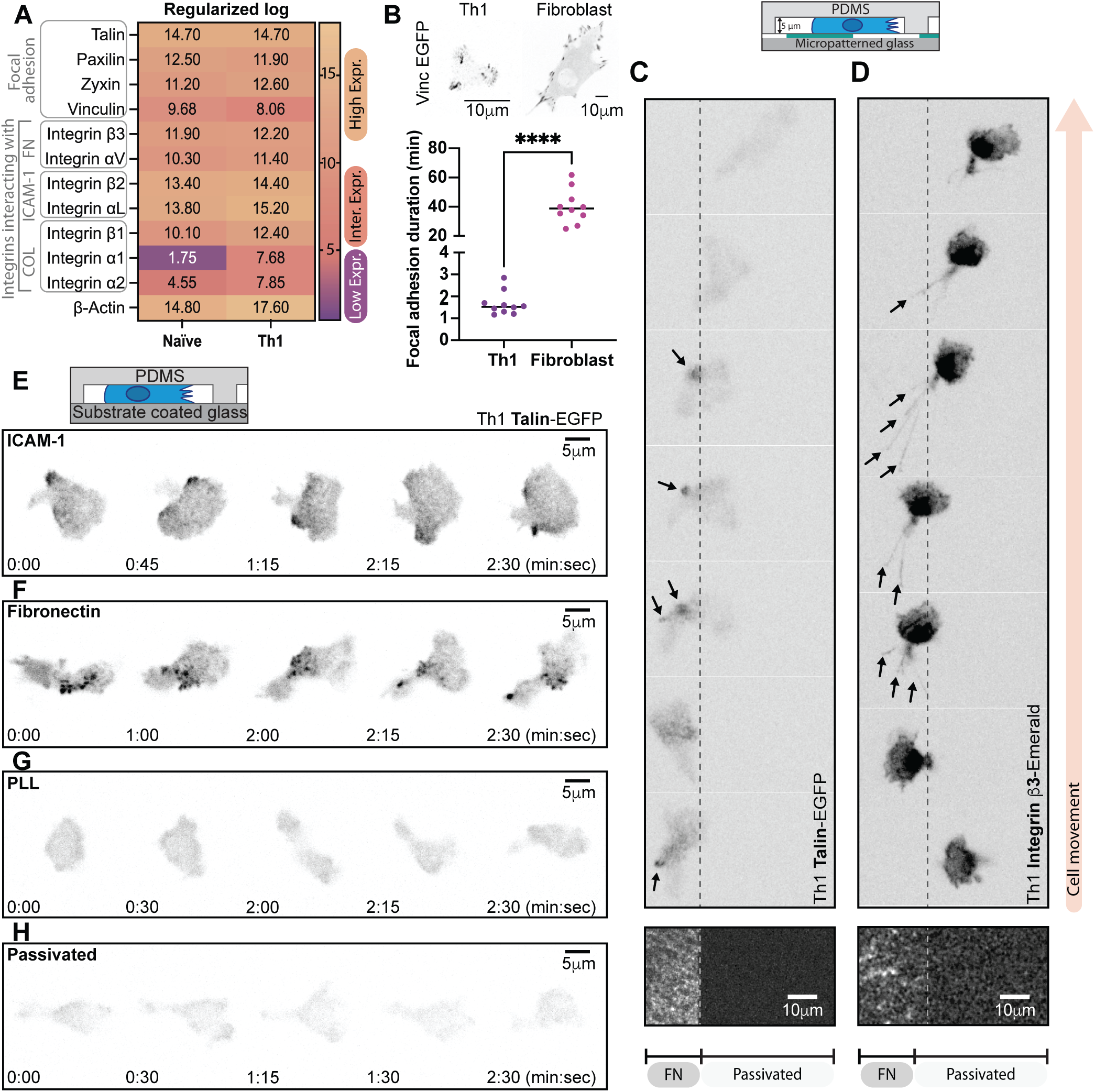
Th1 cells form FAs. (A) RNA-sequencing comparison of naive versus activated T cells (Th1) from publicly available dataset (Th-express.org [43]). Genes shown include FA related proteins, fibronectin-binding integrins, ICAM-1-binding integrins as well as collagen-binding integrins. Their relative levels of expression were calculated from regularised-log (rlog) transformed gene expression counts for all genes and presented using a colormap to show from high, intermediate to low levels of expression. (B) A comparison of FA lifetime between Th1 cells and fibroblasts. Representative Th1 cells expressing Talin-EGFP (C) or Integrin *β*3-Emerald (D) in confinement on a micropatterned substrate. The left portion of the field of view is FN coated and the right portion is passivated. Representative migrating Th1 cells expressing Talin-EGFP in confinement on (E) ICAM-1, (F) Fibronectin, (G) PLL and (H) PLL-PEG.

Finally, to confirm that the FAs were specific to the presence of ECM we imaged cells expressing either talin-EGFP (Fig. 3C, Supplementary Video 6), Integrin β_3_-Emerald (Fig. 3D, Supplementary Video 6) or vinculin-EGFP (Fig. S2E) in Th1 cells migrating across boundaries between fibronectin and passivated substrates. In the talin and vinculin transduced cells, FA-like structures could be seen forming on the ECM coated regions and disappearing as the cell moved onto the passivated regions of the substrates (Fig. 3C, Supplementary Video 6 and Fig. S2E). In the integrin β_3_ transduced cells, it was challenging to identify specific FA structures due to the membrane localization of the probe, but distinct retraction fibers could be seen forming on the ECM coated portion of the substrate (Fig. 3D, Supplementary Video 6), demonstrating clear attachment to the ECM [44]. These results reveal that not only do Th1 cells possess all the necessary components to form FAs, but they only form these structure when ECM or adhesive ligands are present.

### Th1 generate forces on the ECM through focal adhesions

Given the appearance of FA structures, complete with the requisite adhesion proteins [45], we next set out to use traction force microscopy (TFM) [46] to determine whether forces were being exerted at these locations. As Th1 cells require confinement to adhere strongly to FN coated surfaces, we either layered low temperature melting agarose on the gel, or confined them between two FN-coated acrylamide gels, and measured. Most notably, the forces exerted by activated Th1 cells were of a much smaller magnitude than the comparable forces in fibroblasts (Fig. 4A-B). To compare the different cell types we measured both the magnitude of traction stresses and the angle between the traction stress direction and the centroid of the cell (Fig. 4C). While traction stresses in the fibroblasts were in the kPa range (Fig. 4D), the magnitude of traction stresses produced by Th1 cells were a full order of magnitude lower in the 100s of Pa range (Fig. 4E). A similarly strong difference was seen in the distribution of the forces with respect to the cell centroid. In fibroblasts the traction stresses are almost completely contractile (i.e skewed heavily towards θ < 90*^◦^* or directed towards the centroid; Fig. 4D), consistent with previous work [46, 47]. In the Th1 cells, however, we saw a distribution of traction stresses across all angles with an enrichment of traction stresses at both extremities of the distribution. This suggests that the forces were mostly split between pointing toward the cell body (contractile forces, θ < 90*^◦^*) and away from the cell body (pushing forces, θ > 90*^◦^*) (Fig. 4E).

**Fig. 4.**
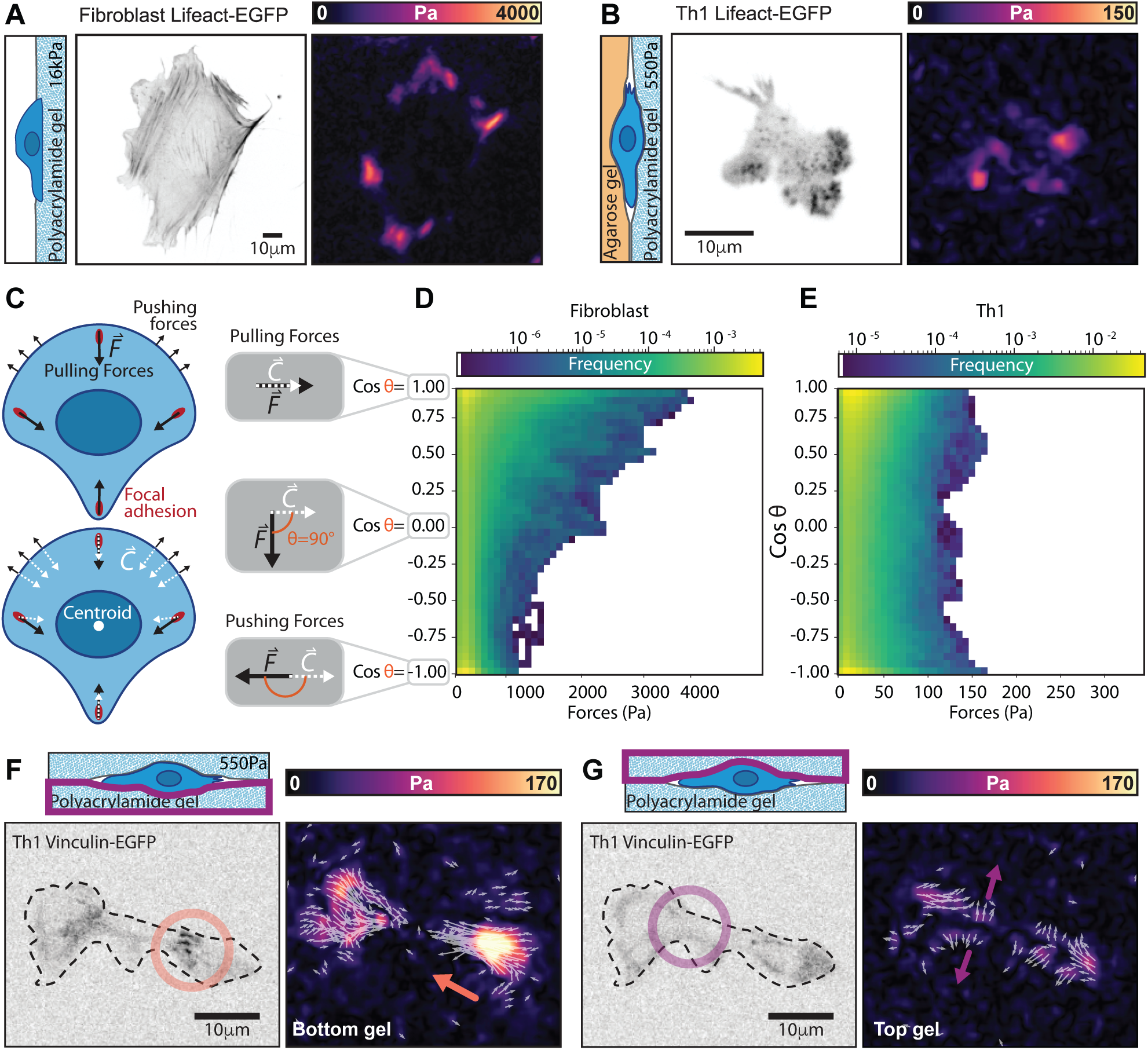
Th1 cells exert traction stresses at FAs. (A-B) Representative traction maps of a (A) fibroblast or (B) Th1 cell expressing Lifeact-EGFP. The fibroblast is plated on a gel with a shear modulus of 16 kPa, while the Th1 cell is plated under agarose on a gel with a shear modulus of 336 Pa. Both gels are coated with FN. (C) Cartoon illustrating the direction of potential forces (black arrows) and their orientation with respect to the cell centroid (white arrows). (D-E) 2D histograms comparing the magnitude of measured traction stresses and the difference in their direction (*θ*) from the cell centroid. Pulling (contractile) forces are represented as cos *θ* = 1, while pushing forces are represented as cos *θ* = *−*1. Histograms are shown for (D) fibroblasts and (E) Th1 cells). (F) Representative images of a Th1 cell expressing Vinculin-EGFP sandwiched between two acrylamide gels of 460 Pa coated with FN. On the bottom gel contractile forces (orange arrow) colocalize with vinculin puncta (orange circle). (G) On the top gel pushing forces (purple arrow) show no colocalization with vinculin (purple circle) and likely arise from the cell pushing the gel out of the way as it is pulled through the gel.

To explore this intriguing finding further we performed double sided traction force microscopy where the Th1 cells were sandwiched between two FN coated polyacrylamide gels to measure how they interacted with their environment. We consistently found that contractile forces coincided with localization of FA proteins such as vinculin (Fig. 4F, Supplementary Video 7), regardless of whether they were on the top or bottom gel. We also observed pushing forces (i.e. θ > 90*^◦^* or directed outward from the cell centroid), that were generally of smaller magnitude and that did not coincide with any visible enrichment of vinculin (Fig. 4G, Supplementary Video 7). In the absence of specific interaction with the substrate, these "pushing" forces likely passively arise in response to the cell body displacing the surrounding gel as the contractile cytoskeletal generated forces exerted at FAs pull the cell forward along its path. The main mechanical interactions driving migration in Th1 cells thus appear very similar to the traditional mesenchymal modes of migration, just at much smaller magnitudes (Fig4A-B) and more rapid timescales (Fig3B).

### ECM promotes confined migration

We next measured migration of activated Th1 T cells confined between a soft acrylamide gel and low-temperature melting agarose. We chose to pour the agar over the cells on the gels, so as not to bias our results by only measuring the cells that could squeeze between the gel and agar. Surprisingly, when the polyacrylamide gels were uncoated, only *≈* 10% of cells were able to migrate at all (Fig. 5A), reinforcing our previous results that Th1 cells require adhesive ligands to migrate (Fig. 2). If the acrylamide gel was coated with either FN or ICAM-1, however, around *≈* 50% of the cells were able to migrate (Fig. 5A). Comparing only the mobile populations of cells on the different ECM coatings, Th1 cells on ICAM-1 migrated significantly further than those on uncoated acrylamide gels (Fig. 5B-C). Mobile cells on FN exhibited a similar trend but with a much larger variance (Fig. 5B-C). These results confirm that ECM is a critical component for migration of Th1 cells through confined environments.

**Fig. 5.**
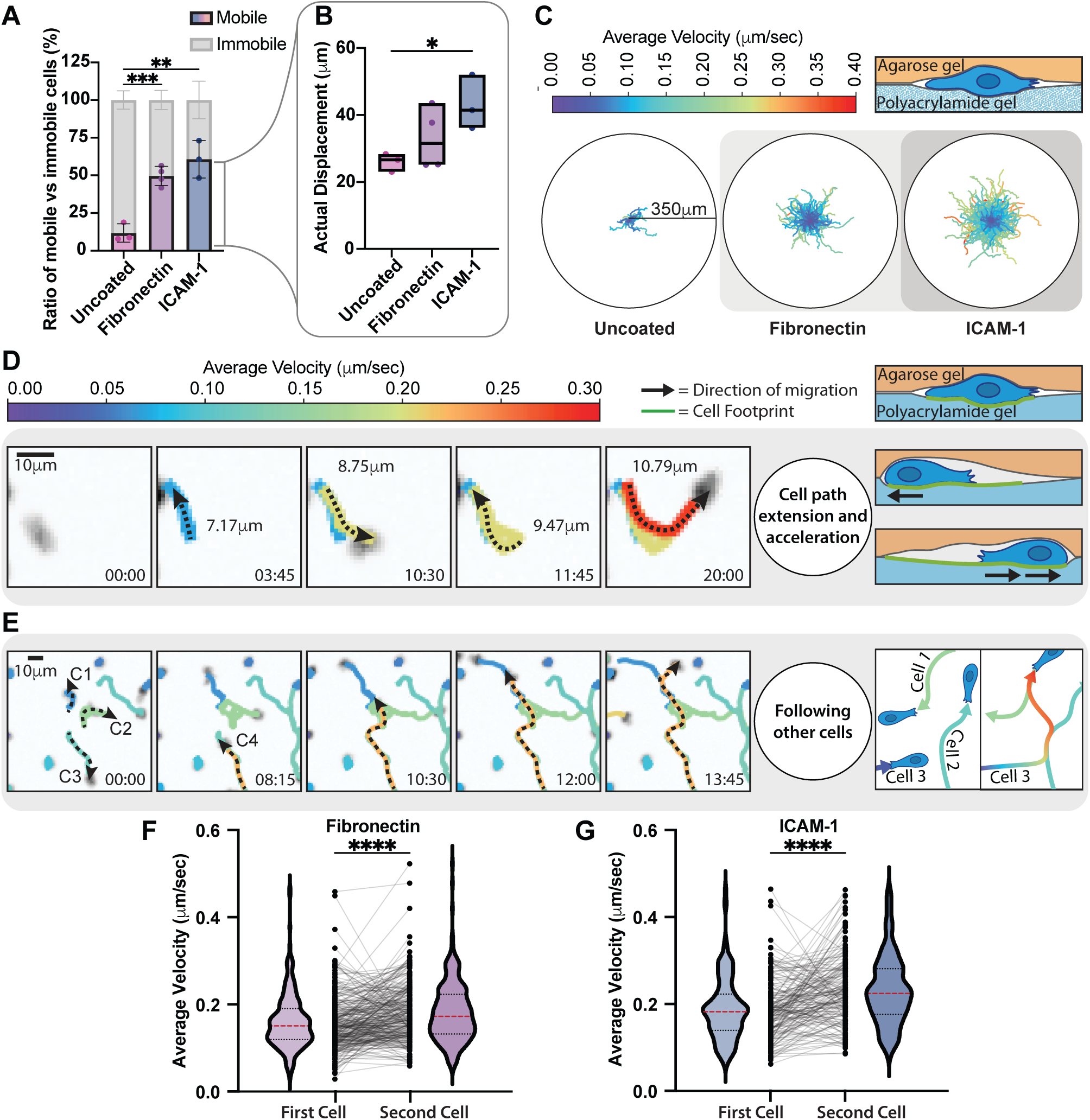
T cells follow the paths of each other. (A) Comparison of the mobile versus immobile fraction of Th1 cells in confinement between an agarose gel and a polyacrylamide gel coated with fibronectin, ICAM-1 or uncoated. (B) Comparison of actual displacement of the cells from (A). Mobile cells are defined by a minimum displacement of 10*µ*m. (B) Roseplots showing Th1 cell tracks of cells in (A, B). The colormap represents the cell average velocity. (D) Individual cell tracking shows cells extending their path and accelerating after each pass. (E) Individual cell tracking shows 3 cells extending their paths and a 4th cell (orange) reusing those previously made paths and accelerating while doing so. Colors represent the cell average velocity. (F-G) Measurement of the change in velocity of cells as they encounter previous paths of migrating T cells confined between an agarose gel and a polyacrylamide gel coated with fibronectin(F) and ICAM-1(G)l.

### FN provides environmental cues to guide migration

It has been previously shown that many immune cells have developed mechanisms to recruit and guide other cells toward inflammation [48]. In our in vitro setup, Th1 cells exhibit similar behaviors, following paths apparently demarcated by previous cells. In these cases we found two interesting observations: First, cells often migrate back and forth along a path, extending it with every pass (Fig. 5D), similar to what has been previously observed in fibroblasts [49]. Second, cells that migrate along a previously trodden path, whether made by themselves or another cell, migrate faster (Fig. 5D-G, Supplementary Video 8). In particular if we compared velocities of cells in regions where trajectories overlapped, we find that on both ICAM-1 and FN the cell passing through second moves faster than the first cell that moves through the same area (Fig. 5F-G). This suggests that cells could potentially be modifying the confined environment either biochemically (e.g. leaving behind a chemokine or degradation of the ECM) or physically (e.g. mechanical deformation of the ECM or membranous tubular network [44]). Either case implicates that FAs and their associated ECM play crucial roles in facilitating migration of Th1 cells through complex environments, beyond simply anchoring the cells to the substrate.

## Discussion

In this paper we demonstrate that activated primary Th1 T cells are able to form FAs and use them to transmit contractile forces generated by the cytoskeleton toward the ECM. We use primary murine helper T cells specifically because they have been shown to increase their expression of β_3_ integrins, which bind to FN, when activated [1, 37], and they are more likely to encounter ECM proteins as they leave the circulatory system to respond to infection. Because of the heterogeneity found in living tissue [50], we chose to perform these experiments in vitro to maximize our control over the extracellular environment. Traditionally, in vitro assays studying T cell migration have relied on coating the substrate with ICAM-1, a ligand which T cells bind strongly. We find that T cells can migrate similarly on FN, an important ECM protein, but only when in confinement. Surprisingly, only a small concentration of FN (such as found in the serum added to media) is required for T cells to migrate. T cells plated integrin independent substrates failed to migrate, but could be rescued simply by coating the substrates with FN or ICAM-1. This interaction with adhesive proteins is facilitated by FAs consisting of the canonical proteins associated with adhesions, including integrins, talin, zyxin, and vinculin, and coincides with regions of contractile forces exerted on the ECM. Furthermore, we find that when migrating, Th1 cells will often follow paths formed by other cells, increasing their migration speed. This follow the leader behavior is also often seen in vivo, suggesting that the T cells may be either modifying the ECM matrix or leaving behind biochemical cues for other T cells to interpret. In total, these results demonstrate that specific cell-substrate interactions are vital components for proper T cell migration, and thus an effective immune response.

The literature has predominantly split different forms of migration into two camps: mesenchymal, encompassing integrin mediated migration, and amoeboid, encompassing everything else [22]. Amoeboid migration derives from the study of the seemingly non-specific interactions between the cell membrane protrusions and the substrate of amoeba such as dictyostelium [51, 52]. Over the years, additional non-specific migration mechanisms, including bleb-based motility [53, 54], pressure driven gradients [55], and even swimming [56] have often been included in the amoeboid category [57]. Migrating immune cells, including T cells, display rapid protrusions, morphology changes, and strong actin flows, consistent with descriptions of amoeboid migration, while lacking the big stable adhesion plaques found in mesenchymal cells [58–60]. This narrative has been furthered by findings that dendritic cells can migrate in 3D following knock-out of all integrins [29], and that both dendritic cells and T cells can migrate on passivated surfaces when in confinement [30, 31]. Recent work, however, has shown that many immune cells do make focal adhesion structures in different contexts [33, 34, 61–64], and T cells specifically form integrin-mediated adhesion when migrating on the planar surface of blood vessels [65–67] and similar structure at the immunological synapse [68–70]. These results suggest that immune cells have the ability to use a variety of mechanisms to migrate in response to different environmental conditions.

With their rapid turnover and small size, the FAs in Th1 cells resemble nascent adhesions seen in mesenchymal cells [71, 72]. The lack of bundled actin stress fibers in T cells facilitates the rapid turnover and short lifetimes of FAs, as actin bundling helps to stabilize these structures in mesenchymal cells [72, 73]. While T cells can form FAs in 2D on ICAM-1 coated surfaces, confinement is required for T cells to migrate on FN coated substrates. The act of confining cells likely increases the number of integrins interacting with the ECM at any given point, helping to stabilize the entire FA as seen in other systems [74–76]. Integrins, including both α*_v_*β_3_ [77] and α*_L_*β_2_ [78], also behave as catch-bonds, with their bond lifetime increasing with load. Confinement may thus increase the interaction of the cell with the substrate, thereby increasing the force of retrograde flow in the cortex and strengthening integrin-ligand bonds. In the tissue, changes in the stiffness of the ECM which are typically associated with inflammation [79], could also increase the load across the bonds, making them last longer.

Strikingly, only a small amount of ECM is required to induce T cell migration. While cells can migrate on passivated surfaces in the presence of serum, migration was significantly reduced when the serum was removed, but could be rescued by coating the substrate with FN. Similarly, when T cells were sandwiched between two uncoated acrylamide gels, the vast majority of T cells were unable to migrate. Previous studies have shown that even when passivated, ECM proteins can still adsorb to surfaces [40]. Together, these findings suggest that passivated glass and PDMS substrates likely contain a small amount of ECM when there is serum present, and this small amount is sufficient to enable migration. Our data also highlights that the percentage of cells that migrate in a given condition is highly variable, and that even in incredibly challenging conditions, some T cells are still able to migrate. This is first and foremost a testament to the robustness of the immune system, but also illustrates the large variation in motility across the population and the importance of avoiding bias by only measuring motile cells.

That we see both specific pulling forces at FAs and non-specific pushing forces on the surrounding ECM during migration is consistent with the idea that ECM geometry strongly influences immune cell migration [80–82]. Adhesions like we see here would be a natural mechanism to sense these different physical properties [50, 83], and could be involved in modifying the external environment to allow following cells to move faster. Indeed, such a pathway was recently proposed with modeling results suggesting that secretion or modification of the substrate could strongly bias the paths of migrating cells [84], consistent with what we see in cells that follow the path of a previous cell (Fig. 5D-G).

In this study we focused on activated primary Th1 T cells, on account of their known expression of FN ligands and their increased likelihood to encounter ECM protein in tissues as they migrate during an immune response. We believe it likely, however, that other immune cells behave similarly. Importantly, our model doesn’t rule out true integrin independent migration, but rather suggests that, if a cell has the components to make an adhesion and ligands are available, it likely will. Ultimately, the difference between amoeboid and mesenchymal migration is blurry at best, as the mechanisms behind true integrin independent migration remain to be explored. Lastly, it is also likely that these FAs play important additional functional roles, such as leaving behind chemokine trails [48] or guiding other cells [44], and future work will be needed to decipher these important roles.

## Materials & Methods

### Primary T cell activation and culture

The workflow of primary T cell collection and activation is illustrated in Fig. S1A. Spleen and lymph node cells were isolated from OTII mice, that express a T cell receptor that recognizes ovalbumin. The next day, 60-80 million cells were stimulated with antigen under Th1 conditions : 4 million viable cells per mL in DMEM AB+ media (High glucose DMEM (HyClone, SH30243.01) supplemented with 10% FBS (Gibco, 10437.028), 2.5% HEPES solution 1M (Hyclone, SH3023701), 1.3% MEM Nonessential Amino Acids 100x (Fisher Scientific, 11-140-050), 0.1% B-ME 55mM (2-Mercaptoethanol, Gibco, 21895-023), 1.2% L-Glutamine (HyClone, Sh30034.01), 1.3% Antibiotic-Antimycotic (Corning, 30-004-CL), 40µg/mL Anti Murine IL-4 (11B11; NCI-Frederick, 05060201), 10 units/mL IL-2 (NCI-Frederick, Teceleukin Recombinant Human Interleukin-2 (rIL-2) - Bulk Ro 23-6019), 2µg/mL OVA peptide (Biomatik, OVA 323-339 custom order), 20ng/mL IL-12 p70 (Peprotech, Cat# 210-12). 48h later viable T cells were isolated on a Lymphocyte Separation Media (LSM) gradient (Corning, 25-072-CV) and either incubated for cells expansion or transduced with retrovirus (see below). After initial activation, the purity of CD4 T cells expressing the OTII T cell receptor was confirmed by flow cytometry (BD LSRFortessa Cell Analyzer) after staining with Anti-CD4-Alexa647(BD bioscience,557681) and Anti-Vα2-PE(BD bioscience,553289). Typically, 30-40% of cells were CD4/Vα2 double positive on day 3 (Fig. S1C) and 70-90% on day 5-9 (Fig. S1D-F).

### Retroviral productions and infection

The empty backbone MIGR1 (Addgene #27490) was used to make retroviral constructs to express Lifeact-EGFP, Integrin β_3_-Emerald, Vinculin-EGFP, Talin-EGFP. Retroviral DNA constructs were transiently transfected into the ecotropic retroviral packaging line, Phoenix. In some cases, Phoenix cells stably producing ecotropic retrovirus were generated by first producing VSV-G pseudo-typed virus and using that to transduce Phoenix cells. Viral supernatants were collected and concentrated 20x using Retro-X concentraror (Takara Bio, PT5063-2), and stored at -80°C. For T cell transduction, on day 3 of activation (Fig. S1A) 250 µl of concentrated virus was preincubated with 8 µg/mL polybrene (EMD Millipore, TR-1003-G) for 30 min on ice, mixed with 1-4 million density purified T cells (see above), and spun for 60 min at 2000RPM at 4°C in a 24 well plate.

### In vitro Live Cell imaging

Experiments with Th1 cells were done in DMEM AB+ media while fibroblasts were plated in DMEM (see cell culture section). All imaging was performed at 37*^◦^*C with 5% Co2 on an Axio Observer 7 inverted microscope (Zeiss) attached to a W1 Confocal Spinning Disk (Yokogawa) with Mesa field flattening (Intelligent Imaging Innovations), a motorized X,Y stage (ASI), and a Prime 95B sCMOS (Photometrics) camera. Illumination was provided by a TTL triggered multifiber laser launch (Intelligent Imaging Innovations) consisting of 405, 488, 561, and 637 nm lasers, using either a 63X, 1.4 NA Plan Apochromat or 10X 0.45NA Plan Apochromat objectives (Zeiss). Temperature and humidity were maintained using a Bold Line full enclosure incubator (Oko Labs). The microscope was controlled using Slidebook 6 Software (Intelligent Imaging Innovations). All in vitro imaging was performed as single confocal slices. For traction force microscopy experiment, FA proteins and the gel were imaged at the same focal plane. Cell were imaged for 20 min at 15 seconds intervals. When cells were imaged in confinement, confinement was applied to the cells immediately before mounting the cell chamber on the microscope in a humidified and tempered chamber, then allowed to stabilize for 5-10 min prior to commencing imaging.

### Cell confinement experiments

Cells in Figs. 1,2 and3 were confined under a PDMS surface with a gap of 5µm as previously described [85, 86]. Briefly, we used SU8 as mold (courtesy of Piel Laboratory, Insitut Curie) to make a confinement slide (CS) that was covered with 5 µm height pillars. Cells were plated on glass coverslip in a cell chamber. A big PDMS pillar attached to a magnetic lid was used to push down the CS onto the cells. The bottom coverslip of the CS were both coated prior to an experiment (see Surface coatings section). In Figs. 4 and 5 we confined cells by using a soft acrylamide gel and either 1% Low Melting Agarose (Invitrogen, 16520-100) or a second soft acrylamide gel. The protocol for soft polyacrylamide gel preparation is described in the section Traction Forces Microscopy Experiments. 1 million cells in 1 mL of DMEM AB+ with 1:10000 Hoechst 33342 (Invitrogen, H3570) were plated on a soft polyacrylamide gel in a coverslip cell chamber and incubated for 10 minutes at 37*^◦^*C or until the cells settled down. The media was removed and 1mL of soft agarose (1% Low Melting Agarose (Invitrogen, 16520-100) in DMEM AB+) previously tempered at 37*^◦^*C was poured on top of the cells. The cell chamber was kept at 37*^◦^*C for 10-20 min to let the cells settle down again while the agarose gel was still unpolymerized. Right before imaging the cell chamber was placed on the ventilation of the tissue culture hood to help reduce the temperature faster and allow the agarose to polymerize for 5 min. Finally 1 mL of DMEM AB+ was added on top of the agarose gel.

### Surface coatings

For FN, glass coverslips were coated with 10 µg/mL human plasma fibronectin (FC010, Millipore) overnight at 37*^◦^*C. For ICAM-1, glass coverslips were first coated with 10 µg/mL Recombinant Protein A (Novex, 10-110-0) overnight at 37*^◦^*C, then blocked with 2% BSA (Bovine Serum Albumin, Research Products International, 9048-46-8) 30 min at RT, then coated with 5 µg/mL ICAM-1 (Sino-Biological, 50440-M03H) 2hr at RT, and finally blocked with 2% BSA (Bovine Serum Albumin, Research Products International, 9048-46-8) 30 min at RT. For PLL coated substrates glass was coated with 0.01% PLL (Poly-L-Lysine, Sigma, P4707) for 20-45 min at RT . For passivated substrates glass was coated with either 1 mg/mL PLL-PEG (JenKem Technology, PLL20K-G35-PEG2K) or 1mg/mL PMOXA (SuSoS, PAcrAm™-g-(PMOXA, amine, silane)) for 20-45 min at RT. For each coatings the coverslips were rinced and kept in PBS at 4*^◦^*C until used (same day for PLL and Passivated coverslips, same or next day for FN and ICAM-1). For polyacrylamide gels after polymerization, gels were rehydrated overnight in ddH2O, treated with cross-linker Sulfo-Sanpah (22589, Pierce Scientific) and photoactivated for 5 min, then washed in ddH2O. Polyacrylamide gels were then immediatly coupled to either human plasma fibronectin (1 mg/mL overnight at 37*^◦^*C; FC010, Millipore) or ICAM-1 (0.1 mg/mL Recombinant Protein A (Novex, 10-110-0) overnight at 37*^◦^*C, 2% BSA (Research Products International, 9048-46-8) 30min at RT, 37.5 µg/mL ICAM-1 (Sino-Biological, 50440-M03H) 2hr at RT, 2% BSA (Research Products International, 9048-46-8) 30 min at RT) .

### Micropatterning experiments

Desired micropattern were designed in AutoCAD and sent to the Nanotechnology Core Facility of University of Illinois Chicago (https://ncf.uic.edu) to make a quartz chrome mask. Before usage the mask is thoroughly washed with water, then isopropanol, and then air dried with an air gun. Glass coverslips were treated with plasma (Plasma Cleaner, Harrick Plasma, PDC32G) for 2 min, then coated with PMOXA (SuSoS, PAcrAm™-g-(PMOXA, amine, silane)) at 1 mg/mL for 45 min at room temperature, then rinsed in ddH2O. The coverslip was placed on the mask, the PMOXA coated side facing the chrome side of the mask. A piece of glass, with similar dimensions to the mask, is then placed on top of the coverslips to hold it in place. The mask-coverslip-glass sandwich is then held together in a custom magnetic frame. The sandwich was then placed in a preheated UVO cleaner oven (Jelight, Model 342), quartz side up, and exposed to deep UV for 4 min. The coverslips were then rinsed in water and coated with 20 µg/mL fibronectin (1/10 Rhodamine Fibronectin (Cytoskeleton, FNR01), 9/10 fibronectin (FC010, Millipore) for 30 min at RT. Micropatterned coverslips were placed in coverslip’s cell chamber, with 250k-500k cells. Cells were then allowed to settle for 10 min before imaging or before confinement.

### Focal adhesion turnover analysis

FA turnover was measured in minimum 12 individual cells for each cell type (Th1 and Fibroblast). For each individual cell the turnover of 5 focal adhesions was analyzed per cell. To analyze turnover, a region of interest (ROI) was drawn around an FA at it’s largest size. We manually tracked the FA over time starting one frame before it appeared until its complete disappearance and recorded that length of time as the FA lifetime.

### Western blot

Protein samples were prepared using 4 million Th1 cells, spun down, rinsed with 1X PBS (Corning, 21-040-CV) and extracted in 500 µL 1X Laemmli Sample Buffer (Bio-Rad, 1610737). The samples were then boiled at 90*^◦^*C for 5 min. The protein quantity was measured using RC DC protein assay Kit II (Bio-Rad, 5000121EDU). For each protein of interest 10 µg of sample was loaded in a 4-15% Tris-Glycine gel (Bio-Rad, 4568083). The gel was transfered onto PVDF membrane (GenScript, L00726) using Genscript eBlot (GenScript, L00686) for 16 min. The membrane was air dried then blocked using 5% nonfat dry milk (Bio-Rad, 1706404) in 1X PBS (Corning, 21-040-CV) with 0.1% Tween 20 (Fisher Scientific, BP337-500) (PBS-T) for 30 min, then rinsed 3X 5 min with PBS-T. The PVDF membranes were incubated overnight at 4*^◦^*C in 2% BSA (Research Products International, 9048-46-8) with primary antibody ; 1:1000 Anti-Talin 1 (Abcam,ab157808), 1:1000 Anti-Zyxin (Millipore Sigma, ABC1463), 1:500 Anti-Paxillin, clone 5H11 (Millipore Sigma, 05-417), 1:1000 Integrin β2 (Cell Signaling, 47598) and 1:1000 Integrin β3 (Cell Signaling, 13166). They were then rinsed 3X 5 min with PBS-T before incubation for 30min in 5% non fat dry milk in PBS-T with secondary antibody ; 1:5000 anti-Mouse HRP antibody (Bio-rad,5178-2504) and 1:5000 anti-Rabbit HRP antibody (Bio-rad,5196-2504), then washed 3X 5 min with PBS-T. Finally the membranes were revealed with Clarity ECL (Bio-Rad, 1705061) for Talin, Zyxin, Paxillin, Integrin β3 and Clarity-Max ECL (Bio-Rad, 1705062) for Integrin β2 using a ChemiDoc (Bio-Rad) imaging system.

### Traction Force Microscopy experiments

Traction force microscopy was performed as described previously [46, 87]. Coverslips were prepared by incubating with a 2% solution of (3-aminopropyl)trimethyoxysilane (313255000, Acros Organics) diluted in isopropanol. Coverslips were washed with DI water 5X for 10 min and cured overnight at 37*^◦^*C. Coverslips were incubated with 1% glutaraldehyde (16360, Electron Microscopy Sciences) in ddH20 for 30 min at room temperature and washed 3X for 10 min in distilled water, air dried and stored at RT. Polyacrylamide gels with a shear modulus of 336 Pa or 460 Pa (for Th1 cells) and 16 kPa (for fibroblast) were embedded with 0.04-µm fluorescent microspheres (F8789, Invitrogen) and polymerized on activated glass coverslips for 30 min - 1 h at room temperature. After polymerization, gels were rehydrated overnight in ddH2O, treated with cross-linker Sulfo-Sanpah (22589, Pierce Scientific) and photoactivated for 5 min, then washed in ddH2O. Polyacrylamide gels were then immediatly coupled to matrix proteins, human plasma fibronectin (1mg/mL overnight at 37*^◦^*C; FC010, Millipore) or adhesion protein, ICAM-1 (0.1 mg/mL Recombinant Protein A (Novex, 10-110-0) overnight at 37*^◦^*C, 2% BSA (Research Products International, 9048-46-8) 30min at RT, 37.5 µg/mL ICAM-1 (Sino-Biological, 50440-M03H) for 2hr at RT, 2% BSA (Research Products International, 9048-46-8) for 30m in at RT) . Following matrix protein cross-linking, gels rinsed in PBS and kept at 4*^◦^*C in PBS until used (either same day or next day). For fibroblast cells, they were plated on the gels, allowed to adhere overnight and imaged the following day. For Th1 cells, they were plated on the gels 10-20 min before confinement by low melting agarose or a second polyacrylamide gel. Reference image of unstrained gel was obtained by waiting for the cell to migrate out of the field of view.

For 336 Pa gels, a standard solution was prepared with 1.25 mL of 40% Acrylamide (Biorad, 1610140) 583 µL 2% bis-Acrylamide (Biorad, 1610142) and 3.16 mL ddH2O. To make the gel 150µL of standard solution was mixed with 341.75 µL water, 5 µL 0.04-µm fluorescent microspheres (F8789, Invitrogen), 0.75 µL TEMED (Fisher Bioregents, 110-18-9) 2.5µL of 10% APS (Ammonium Persulfate, Fisher Bioregents, BP179-25). 7 µL of the mixture was used for a 22x30 mm coverslip.

For 460 Pa gels, a standard solution was prepared with 1.25 mL of 40% Acrylamide (Biorad, 1610140) 666 µL 2% bis-Acrylamide (Biorad, 1610142) and 3.08 mL ddH2O. To make the gel 150µL of standard solution was mixed with 341.75 µL water, 5 µL 0.04-µm fluorescent microspheres (F8789, Invitrogen), 0.75 µL TEMED (Fisher Bioregents, 110-18-9) 2.5µL of 10% APS (Ammonium Persulfate, Fisher Bioregents, BP179-25). 7 µL of the mixture was used for a 22x30 mm coverslip.

For 16 kPa gels, a standard solution was prepared with 2.5 mL of 40% Acrylamide (Biorad, 1610140), 604 µL 2% bis-Acrylamide (Biorad, 1610142) and 1.896 mL ddH2O. To make the gel 300 µL of standard solution was mixed with 191.75 µL water, 5 µL 0.04-µm fluorescent microspheres (F8789, Invitrogen), 0.75 µL TEMED (Fisher Bioregents, 110-18-9) 2.5 µL 10% APS (Ammonium Persulfate, Fisher Bioregents, BP179-25). 7 µL of the mixture was used for a 22x30 mm coverslip.

Analysis of traction forces was performed using code written in Python according to previously described approaches [46, 87, 88]. Code is available at https://github.com/OakesLab/TFM. Prior to processing, images were flat-field corrected and aligned to the reference bead image with the cell detached. Other acquired channels were shifted using the same alignment measurements from the bead channel. Displacements in the beads were calculated using an optical flow algorithm in OpenCV (Open Source Computer Vision Library, https://github/itseez/opencv) with a window size of 8 pixels. Traction stresses were calculated using the Fourier Transform Traction Cytometry (FTTC) approach [46, 89] as previously described, with a regularization parameter of 8.87*×*10*^−^*^5^.

For the 2D histograms (Fig. 4D,E) we calculated the centroid of a binary cell mask, and then for each pixel in the cell mask we calculated the vector that pointed from that pixel towards the centroid. If 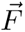 represents the traction stress vector at a pixel and 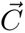 represents the vector towards the centroid, we used the dot product of two vectors to calculate 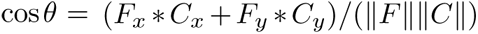. The magnitude of the traction stress was calculated as 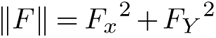. The 2D histogram was then constructed from these two values on a per pixel basis using all pixels contained in the cell masks across all the cells measured in the dataset.

### Gel Stiffness Measurements

Gels were fabricated as described above, with the only difference that a spacer was used during polymerization to create a thicker gel of *≈* 300-350 µm in height. Gel stiffness was measured be measuring the deformation caused by a stainless steel ball bearing 1.5 mm in diameter (Fig. S3A), as previously described [90]. Briefly, the gel height was measured by taking the difference between the bottom and top of the gel. A confocal z-stack with a step size of 1.25 µm was then taken through the top of the gel (Fig. S3C-D), and the deformation was determined by finding the center of the indentation (Fig. S3B)and fitting a circle with radius equivalent to the bearing (Fig. S3E-F). This depth measurement was repeated in two orthogonal directions and averaged (Fig. S3C-F). The gel Young’s modulus was then determined using a modified Hertz model [90, 91] to account for the gel being thin gels bonded to a surface. At least two measurements were taken per gel, with at least two gels per replicate and experiments were repeated in triplicate (Fig. S3G). Gel stiffness value represents the shear modulus.

### Cell migration tracking

All cell tracking was performed using trackpy [92]. Tracks were only included if they lasted more than 5 frames and tracked cells were not allowed to skip frames. Once the track positions were calculated, track parameters were calculated as follows: instantaneous velocity was the difference between positions between frames; path duration was the total number of frames multiplied by the frame interval; path length was the sum of the displacements between each point along the track; the average velocity was the path length divided by the path duration; the actual displacement was the geometric distance between the first and last points of the path; the effective velocity was the actual displacement divided by the path duration. Cells were classified as immobile if the actual displacement was less than 10 µm (Fig.2F).

To determine overlapping tracks, we first made binary images of each isolated track and then computed a sum projection of the binary image stack. Pixels with values greater than 1 indicated regions where trajectories overlapped. The tracks that passed through each of these regions were identified and only the first two cells that passed through a given overlap were analyzed. Each potential overlap of trajectories was visually inspected to ensure that the tracks in fact overlapped. Average velocity measurements were then taken and compared for each cell passing through the overlap ROI, only considering those points that were actually in the overlap.

### Statistical analysis

For all analysis, statistically different results are defined as follow : p-value < 0.05 = *; p-value < 0.005 = **; p-value < 0.0005 = ***; p-value < 0.0001 = ****. Non-statistically different results (p-value > 0.05) are either identified with "ns" or without a star. For all analysis, cells that moved less than 10 µm were considered immobile. All comparisons were made using unpaired parametric t tests, except for Figs. 1B, 5F-G, & S1G-H which used paired parametric t tests. For each replicate, unless otherwise specified, between 1000-2500 cells were analyzed spread across *≈* 5 of fields of view per experiment and the data point represents the mean of the value plotted. The number of experimental replicates for the data presented in the figures is detailed below:

- Fig. 1B: FN, n=3; ICAM-1, n= 4.
- Fig. 1D-F: FN, n=8; ICAM-1, n= 6; PLL, n=6; Passivated, n= 7.
- Fig. 2A-C: PLL n=3; Passivated n=4; 0.1 µg/mL FN n=3; 1 µg/mL FN n=3; 10 µg/mL FN n=4.
- Fig. 3B: Th1, n=10; Fibroblast, n=10. Each replicate compared the turnover of at least 5 FAs per cell
- Fig. 4D-E: Th1, n=45 from 4 different experiments; Fibroblast, n=12. The data for the histogram was pooled from all cells resulting in *≈* 6 *×* 10^5^ vectors for the Th1 cells, and 2 *×* 10^6^ vectors for the fibroblasts.
- Fig. 5A-B: Uncoated, n=3; FN, n=4; ICAM-1, n= 3.
- Fig. 5F-G: FN, n=4; ICAM-1, n= 3. The paired data was pooled and consisted of 268 total overlaps examined on FN and 172 overlaps on ICAM-1
- Fig. S1G-H. All conditions n=3. Samples with and without FBS were performed with the same preparation of cells, on the same day, from the same mice.

### RNAseq data analysis

We use RNA-seq data from the publicly available database (Th-express.org [43]) to compare a selection of different proteins involved in focal adhesion and integrins. The gene name of each of the selected proteins was searched on Th-express.org database. The regularised-log (rlog) measure was collected for each gene. Rlog represents transformed gene expression counts for all genes. The results were presented using a colormap to show from high expression, intermediate expression, and low expression level based on the description of gene frequency in the database.

### Code

All code used to analyze data is available at https://github.com/OakesLab.

## Author Contributions

AC, DJF, PWO conceived and designed the study. AC performed all experiments and made the figures. DO and JM developed constructs, handled animals and prepared tissue for T cell isolation. AC and PWO performed data analysis. AC and PWO wrote the manuscript with input from all other authors.

## Acknowledgements

We want to thank all the members the Beach and Oakes laboratories at Loyola, the Topham and Kim labs at the University of Rochester, and the Fowell lab at Cornell for their helpful feedback. We also want to thank the Piel lab from Institut Curie for their help in the confinement techniques, and Dr. Stefano Sala for assistance with the gel stiffness measurements. A.C acknowledges support from the Company of Biologist for financing a training visit to the Piel lab. P.W.O. acknowledges support in part by a National Science Foundation CAREER Award #2000554. D.J.F., J.M. and P.W.O. acknowledge support from the National Institutes of Health (NIH) National Institute of Allergy and Infectious Disease (NIAID) Award P01-AI102851.

## Declarations of Interest

None

## Notes

### Competing Interest Statement

The authors have declared no competing interest.

