## Supplemental Figures for "T cells Use Focal Adhesions to Pull Themselves Through Confined Environments"

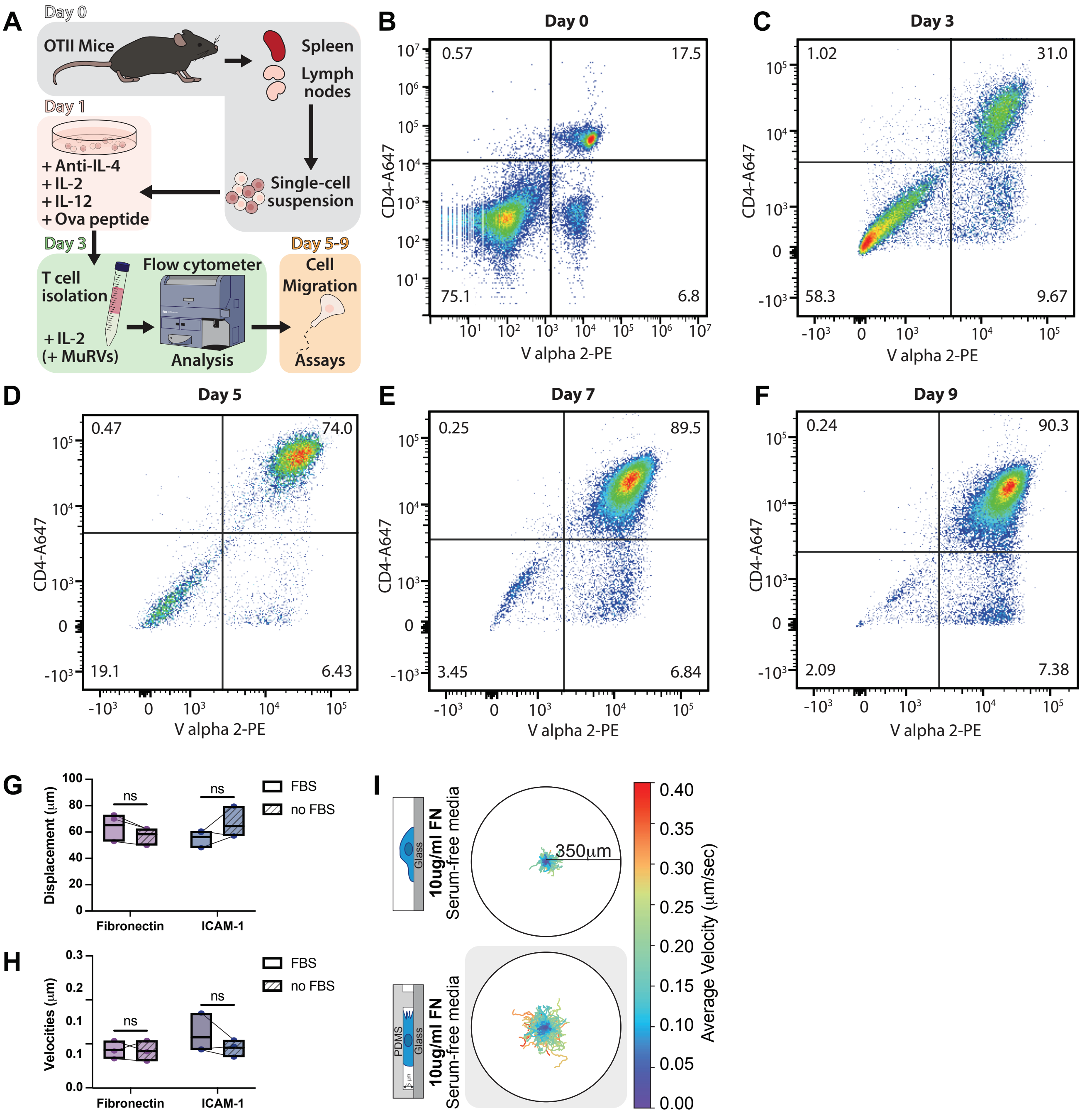


**Fig. S1.** (A) Graphical representation of Th1 activation. (B) Flow cytometer analysis of a representative single cell suspension on the day of extraction from OTII mice using CD4 and V alpha 2 antibodies as markers. (C) A representative flow cytometer analysis of T cells 3 days post activation with anti-IL-4 (11B11; 40ug/ml), IL-2 (20units/ml), IL-12 (40ng/ml), OVA peptide (4ug/ml). T cells were isolated using lymphocyte separation media. (D-F) Representative flow cytometer analyses of Th1 T cells after 5 (D), 7 (E) and 9 (F) days post activation, with continuous treatment treatment of 10 units/ml IL-2 and lymphocyte separation media before each use. The actual displacement (G) and effective velocities (H) of mobile cells on fibronectin and ICAM-1 substrate with and without FBS in cell media. (I) Th1 cell tracking in serum-free media on Fibronectin in unconfined vs confined (5$\mu$m) environments. The colormap represents the cell average velocity in $\mu$m/sec.

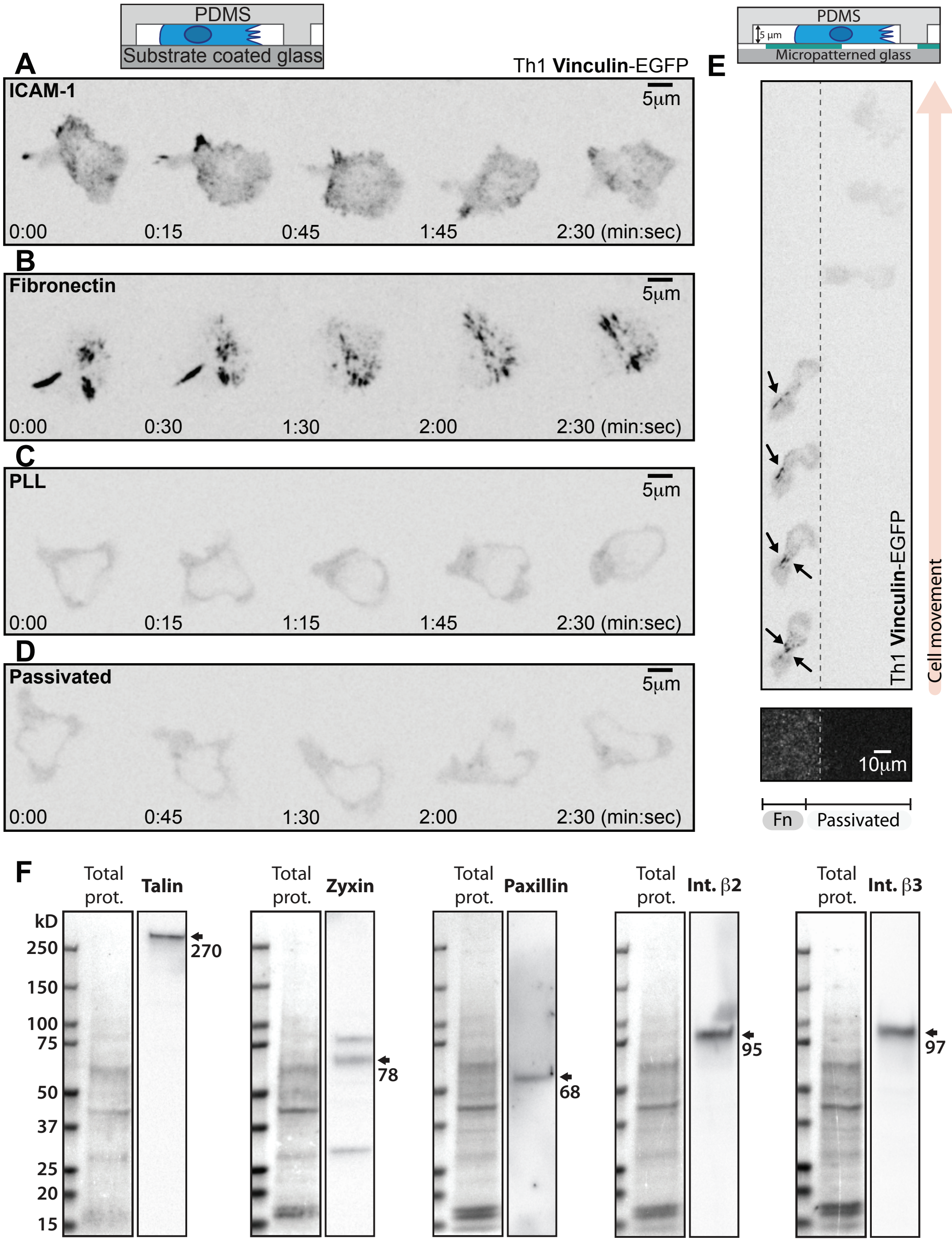


**Fig. S2.** Representative Th1 cells expressing Vinculin-eGFP in confinement on (A) ICAM-1, (B) Fibronectin, (C) PLL and (D) PLL-PEG. (E) A Th1 expressing Vinculin-eGFP in confinement on a micropatterned substrate. The left portion of the field of view is fibronectin coated and the right portion is passivated. Arrows indicate vinculin accumulation in FAs. (F) Representative Western blots showing endogenous expression of FA proteins in activated Th1 cells.

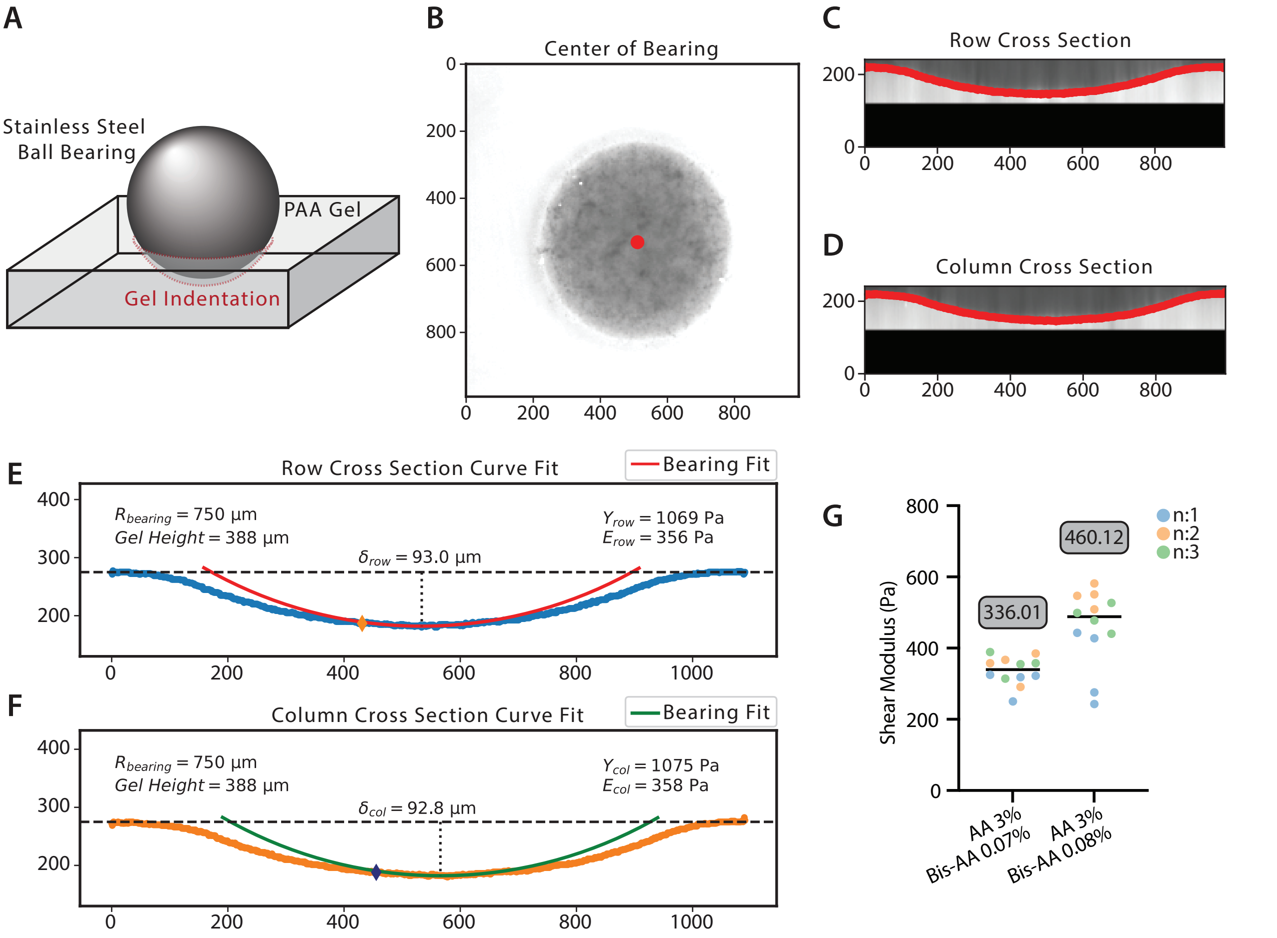


**Fig. S3.** Gel stiffness measurements. (A) Graphical representation of the measurement set up (See Methods for details)(B) Determination of the center of the indentation. (C-D) Orthogonal views of the gel Z-stack passing through the center of the indentation. (E-F) Ball bearing curve fitted to the gel indentation to determine gel stiffness. (G) Representation of 3 biological replicates and 12 technical replicates to determine the average stiffness of each gel recipe used in this manuscript.

### Supplemental Movies

**Supplemental Movie 1.** Th1 cells stained with hoescht migrating in confined (5 $\mu$m) environments coated with ICAM-1, Fibronectin, PLL or Passivated. 10x objective. Frame rate=15 seconds. Movie from Fig.1D-F.

**Supplemental Movie 2.** Left pannel: Th1 cells stained with hoescht migrating on a FN micropatterned substrate (Light gray=FN, Darker gray=Passivated). Right pannel: Cell tracking of the right pannel. The colormap represents the cell average velocity in $\mu$m/sec. 10x objective. Frame rate=30 seconds. Movie from Fig.1H-J.

**Supplemental Movie 3.** Cell tracking of Th1 cells stained with hoescht migrating in confined (5 $\mu$m) environments coated with Fibronectin or Passivated in serum-free media (-FBS) or FBS supplemented media (+FBS). 10x objective. Frame rate=15 seconds. Movie from Fig.2D

**Supplemental Movie 4.** Cell tracking of Th1 Lifeact-EGFP cells migrating in confined (5 $\mu$m) environments coated with Fibronectin or Passivated in serum-free media (-FBS). 63x objective. Frame rate=15 seconds. Movie from Fig.2E

**Supplemental Movie 5.** Th1 Talin-EGFP cells migrating in confined (5 $\mu$m) environments coated with ICAM-1, Fibronectin, PLL or Passivated. 63x objective. Frame rate=15 seconds. Movie from Fig.3E-H

**Supplemental Movie 6.** Th1 Talin-EGFP (Left pannel) and Th1 Integrin $\beta$3-Emerald (Right pannel) cells migrating in confined (5 $\mu$m) environments on a micropatterned substrate. 20x objective. Frame rate=15 seconds. Movie from Fig.3C-D

**Supplemental Movie 7.** Top pannels : Th1 Vinculin-EGFP cells sandwiched between two acrylamide gels of 460 Pa coated with FN. Bottom pannels : Representation of traction stress vector overlay on traction map. Pulling traction stress colocalize with vinculin puncta. Pushing forces show no colocalization with vinculin and seems to outline cell shape. 63x objective. Frame rate=15 seconds. Movie from Fig.4F-G

**Supplemental Movie 8.** Cell tracking of Th1 cells stained with hoescht migrating in confinement between an agarose gel and a polyacrylamide gel coated with fibronectin. The movie shows 3 cells extending their paths and a 4th cell (orange) reusing those previously made paths and accelerating while doing so. The colormap represents the cell average velocity in $\mu$m/sec.10x objective. Frame rate=15 seconds. Movie from Fig.5E
